# *Staphyloccocus aureus* biofilm, in the absence of planktonic bacteria, produces factors that activate counterbalancing inflammatory and immune-suppressive genes in human monocytes

**DOI:** 10.1101/2023.10.06.561208

**Authors:** Richard D Bell, E. Abrefi Cann, Bikash Mishra, Melanie Valencia, Qiong Zhang, Xu Yang, Alberto Carli, Mathias Bostrom, Lionel Ivashkiv

## Abstract

*Staphyloccocus aureus* (*S. aureus*) is a major bacterial pathogen in orthopedic periprosthetic joint infection (PJI). *S. aureus* forms biofilms that promote persistent infection by shielding bacteria from immune cells and inducing an antibiotic-resistant metabolic state. We developed an in vitro system to study *S. aureus* biofilm interactions with primary human monocytes in the absence of planktonic bacteria. In line with previous in vivo data, *S. aureus* biofilm induced expression of inflammatory genes such as *TNF* and *IL1B*, and their anti-inflammatory counter-regulator IL-10. *S. aureus* biofilm also activated expression of PD-1 ligands that suppress T cell function, and of IL-1RA that suppresses differentiation of protective Th17 cells. Gene induction did not require monocyte:biofilm contact and was mediated by a soluble factor(s) produced by biofilm-encased bacteria that was heat resistant and > 3 kD in size. Activation of suppressive genes by biofilm was sensitive to suppression by Jak inhibition. These results support an evolving paradigm that biofilm plays an active role in modulating immune responses, and suggest this occurs via production of a soluble vita-PAMP. Induction of T cell suppressive genes by *S. aureus* biofilm provides insights into mechanisms that suppress T cell immunity in PJI, and suggest that anti-PD-1 therapy that is modeled on immune checkpoint blockade for tumors may be beneficial in PJI.

## Introduction

Periprosthetic joint infection (PJI) is a devastating complication of joint replacement surgery that is resistant to treatment and a major cause of morbidity, and even mortality, in orthopedics (Toulson, Walcott-Sapp et al. 2009, Kurtz, Lau et al. 2012, Berend, Lombardi et al. 2013, Tande and Patel 2014, Kapadia, Berg et al. 2016, Ricciardi, Muthukrishnan et al. 2020). *Staphylococcus aureus* (*S. aureus*) is a major cause of PJI, affecting up to 63% of culture-positive cases (Weiser and Moucha 2015). Lack of clearance of bacterial pathogens like *S. aureus* in response to antibiotic therapy results in persistent infections that cause pain, bone loss, and decreased mobility, necessitating surgical interventions such as debridement with lavage, and challenging revision surgery for implant removal and re-implantation (Costerton, Stewart et al. 1999, Scherr, Heim et al. 2014, Bröker, Mrochen et al. 2016, Goldmann and Medina 2018, Ricciardi, Muthukrishnan et al. 2018, Saeed, McLaren et al. 2019, de Vor, Rooijakkers et al. 2020). One major reason for persistence of PJI is sequestration of microbes in a bacterially-generated biofilm that forms on the implant (Scherr, Heim et al. 2014), and also in bone canalicular network, in abscess communities, and in necrotic tissue (Scherr, Heim et al. 2014, Bröker, Mrochen et al. 2016, Goldmann and Medina 2018, Ricciardi, Muthukrishnan et al. 2018, de Vor, Rooijakkers et al. 2020). An additional key reason for PJI persistence is that pathogens like *S. aureus* can suppress immune cells directly (Miller and Cho 2011, Bröker, Mrochen et al. 2016, Goldmann and Medina 2018, de Vor, Rooijakkers et al. 2020), or indirectly by activation of immune-suppressive pathways and generation of suppressive myeloid cells (also termed myeloid-derived suppressor cells, MDSCs) that suppress adaptive immunity mediated by T cells (Heim, Vidlak et al. 2014, Heim, Vidlak et al. 2015, Tebartz, Horst et al. 2015, Heim, Vidlak et al. 2018, Heim, West et al. 2018, Aldrich, Heim et al. 2020, Schwarz 2022, Sokhi, Xia et al. 2022).

Biofilm consists of a matrix of proteins, extracellular DNA and polysaccharides in which bacteria become embedded and enter a metabolically altered quiescent state in which replication is suppressed by quorum sensing (Yamada and Kielian 2019). One well established mechanism by which biofilm contributes to persistence of infection is physical sequestration of bacteria from cells of the immune system. In addition, the altered metabolic state and replicative quiescence of bacteria within the biofilm renders them resistant to antibiotics. Persistent biofilm-associated infections such as *S. aureus* PJI are often associated with activation of immune-suppressive pathways, infiltration by macrophages polarized towards an M2-like alternatively activated state, infiltration by MDSCs that suppress T cells, and a paucity of T cells, which appear to be in a suppressed state (Miller and Cho 2011, Heim, Vidlak et al. 2014, Heim, Vidlak et al. 2015, Tebartz, Horst et al. 2015, Bröker, Mrochen et al. 2016, Goldmann and Medina 2018, Heim, Vidlak et al. 2018, Heim, West et al. 2018, Yamada and Kielian 2019, Aldrich, Heim et al. 2020, de Vor, Rooijakkers et al. 2020, Schwarz 2022, Sokhi, Xia et al. 2022). This has led to an emerging idea that biofilm and the bacteria contained within can regulate and suppress immune responses. However, biofilm is also a source of planktonic bacteria, which bud from the biofilm surface and drive co-existing planktonic infections in the adjacent tissues that cause tissue damage and pain (Tande and Patel 2014, Ricciardi, Muthukrishnan et al. 2020). Since planktonic phase bacteria such as *S. aureus* also suppress and modulate immune responses, the role of biofilm itself in immune suppression/deviation in PJI and its mechanisms of action are not well understood. Furthermore, biofilm-associated infections also exhibit strong activation of innate immune cells (Krishna and Miller 2012, Menousek, Horn et al. 2022, Sokhi, Xia et al. 2022), and the balance between induction of suppressive and inflammatory pathways by biofilm has not been well studied.

Mechanisms by which biofilm regulates immune cell functions have been investigated using in vitro models, typically using in vitro-generated biofilm and in vitro-differentiated mouse macrophages and myeloid cells (Southey-Pillig, Davies et al. 2005, McBain 2009, Thurlow, Hanke et al. 2011, Hanke, Heim et al. 2013, Wei and Ma 2013, Heim, Vidlak et al. 2014, Scherr, Hanke et al. 2015, Yamada and Kielian 2019). These studies have shown that in addition to controlling nutrient, pH and oxygen gradients that affect immune cell function, bacterial biofilm produces various proteins and small molecules including leucocidin AB and alpha toxin (Scherr, Hanke et al. 2015, Yamada and Kielian 2019) that are either toxic to immune cells or can regulate their differentiation, polarization and metabolism. Live *S. aureus* and biofilm can activate inflammatory genes such as *Tnf* and *Il6* in mouse myeloid cells by epigenetic chromatin-mediated mechanisms (Van Roy, Shi et al. 2023), and bacteria-produced lactate inhibits histone deacetylase 11, thereby enhancing production of the anti-inflammatory cytokine IL-10 in macrophages and MDSCs (Heim, Bosch et al. 2020). In these previous studies, as best as we can determine, the authors did not address the potential role of bacteria that bud from biofilm during in vitro culture, enter a planktonic phase and rapidly replicate during the time course of experiments, and thus can potentially contribute to the observed results.

We had previously found in an established in vivo model of orthopedic *S. aureus* PJI (Carli, Ross et al. 2016, Carli, Bhimani et al. 2017, Sokhi, Xia et al. 2022) that immune cells isolated from the infected implant-bone interface and thus adjacent to biofilm expressed high levels of inflammatory cytokines, comparable to or even higher than infected joint soft tissues or bone more distal to the implant (Sokhi, Xia et al. 2022). We wished to test whether *S. aureus* biofilm, in the absence of planktonic bacteria, activates myeloid lineage cells, whether this is a direct effect related to biofilm:cell contact or mediated by soluble factors, and the balance between induction of inflammatory versus suppressive genes. To maximize relevance for human PJI, we used primary human monocytes that directly correspond to cells that migrate into infected sites and have distinct activation profiles from mouse macrophages. Our work demonstrates that biofilm-produced soluble factors induce a strong and sustained inflammatory gene induction from monocytes as well as counterbalancing induction of immunomodulatory genes. This gene induction is not dependent on direct contact with the biofilm and the immunomodulatory genes can be attenuated by inhibition of Jak kinases.

## Materials and Methods

### Bacterial Strains and Biofilm Culture

*S. aureus* (SA) strain Xen36 containing a kanamycin resistant cassette (PerkinElmer American Type Culture Collection #49525), was obtained (Carli, Bhimani et al. 2017) and used for all in vitro biofilm studies. To generate bacterial colonies, a frozen Xen36 aliquot was expanded in liquid culture overnight and then streaked onto sterile agar plates (3% Tryptic Soy Broth (TSB) with 200ug/mL of kanamycin sulfate) and incubated overnight at 37°C. Individual colonies were then picked and used for all experiments. Individual colonies were inoculated into biofilm culture media (BCM; 4mL of RPMI, 10% Fetal Bovine Serum (FBS), 200ug/mL of kanamycin sulfate and 1% HEPES) and incubated in 15ml round bottom falcon tubes on a shaker at 250 RPM at 37°C overnight. To generate a culture of SA in log phase of growth, 700 µl of the overnight culture was added to 9mL of BCM and incubated in a shaker for ∼1.5 hours until the optical density (OD) of the culture at 600 nm was 0.5 which translates to ∼4×10^8^ colony forming units (CFU). To generate biofilm, 10uL of this culture was plated with 490 ul of BCM in a 24 well cell culture plate and grown at 37°C for 4 days. The media was changed every 24hrs to remove planktonic bacteria while minimally disturbing the biofilm on the bottom of the wells.

### Wheat Germ Agglutinin Assay

Biofilms were grown for 24, 48, or 96 hrs and then washed with PBS (2x), stained with Wheat Germ Agglutinin (WGA) conjugated to FITC (Invitrogen) for 2hrs at RT in the dark, washed again with PBS (3x) and then imaged on a fluorescent microscope (AxioObserver, Zeiss).

### Monocyte and Macrophage Isolation

Deidentified buffy coats purchased from the New York Blood Center (Queens, New York, NY) following a protocol approved by the Hospital for Special Surgery Institutional Review Board. Peripheral blood mononuclear cells were purified from using density gradient centrifugation with Lymphoprep (Accurate Chemical) and monocytes were then isolated using anti-CD14+ magnetic beads (Miltenyi) following the manufacturers recommendations (Yang, Bachu et al. 2022). CD14+ cells were resuspended at 2×10^6^ cells/mL in RPMI containing 10% Fetal Bovine Serum (FBS), 1% HEPES, 20 ng/ml human M-CSF, and 5ug/mL of gentamycin (Monocyte Culture Media, MCM).

### Monocyte/Biofilm Co-Culture and Transwell Experiments

We treated Day 4 biofilm with 50ug/mL gentamycin overnight to eliminate planktonic bacteria, which was verified in pilot experiments by plating biofilm culture supernatants on agar plates and counting colonies. The next day 500 ul of CD14+ cells at 2×10^6^ cells/mL were in RPMI containing 10% Fetal Bovine Serum (FBS), 1% HEPES, and 5ug/mL of gentamycin were added to the biofilm; 5ug/mL of gentamycin was verified in several experiments to prevent outgrowth of planktonic *S. Aureus* from the viable bacteria contained within the biofilm. Control CD14 cells in the same medium were plated in empty wells and treated with 25ng/mL of Pam3cys or left untreated. For RNA extraction, the media was aspirated and RLT buffer (Qiagen) was added to the wells. RNA was then extracted using the Qiagen RNeasy Kit (Qiagen), reverse transcribed into cDNA using RevertAid ET Reverse Transcription Kit (Thermo Scientific), and qPCR was performed using SYBR Green (Applied Biosystems) with the QuantStudio 5 (Applied Biosystems). PCR primers are presented in Supplemental Table 1.

For Transwell (StemCell Technologies) studies, we generated 4-day-old biofilm as described above. On day 5 Transwell inserts were placed into the plates and CD14+ cells were plated in the inserts. Control cells were added into Transwells placed into culture wells that did not contain biofilm.

### Biofilm Supernatant Experiments

To generate biofilm supernatant, we generated 4-day-old biofilm as described above, treated biofilm with 50ug/mL gentamycin for 24 hrs, then cultured with BCM containing 5 ug/ml gentamycin for 12-14 hrs. This supernatant was treated in 6 distinct ways: 1) left unaltered (described as Biofilm Supernatant or Unaltered Supernatant), 2) Heated to 100°C twice for 15 mins allowing the supernatant to cool to RT in between the two heat cycles (Heated Supernatant), 3) centrifuged at 21,000 x *g* for 10 mins then collected and passed through a 0.22µm filter (Spun and 0.22µm Filtered Supernatant), 4) Heated to 100°C twice for 15 mins allowing the supernatant to cool to RT in between the two heat cycles, then centrifuged at 21,000 x *g* for 10 mins and passed through a 0.22µm filter (Heated, Spun and 0.22µm Filtered Supernatant), 5) the supernatant was placed into a 3kDa centrifugation filter (Pierce/Thermo Scientific) and centrifuged at 4000 x *g* for 20 mins, and eluent was collected and used (3kDa filtered supernatant), and 6) Heat inactivated as described above, then centrifuged at 21,000 x *g* for 10 mins then collecting and then passed through a 3kDa centrifugation filter and centrifuged at 4000 x *g* for 20 mins; the eluent was then collected (Heated, Spun, and 3kDa Filtered Supernatant). All supernatants were cultured at a 1:1 dilution with CD14+ cells.

### Colony Forming Unit Assays

Throughout the experiments we performed CFU assays to quantify the amount of bacterial growth in the biofilm cultures. 10 or 50uL of biofilm supernatant was spread on a TSB plate with a L-shaped spreader and incubated overnight and colonies were counted.

### Statistical Analysis

All statistical analysis was performed in Prism 8.0 (GraphPad). All data was tested for normality with a Shapiro Wilks test, and in all cases at least one group was found to be not-normally distributed. Thus, all data was log transformed with the equation log(y+1) and considered normal. Given that within each experiment, all conditions were performed on CD14+ cells from the same donor, we utilized a repeated measures statistical design matching by donor. Two-way ANOVAs with Tukey’s post-hoc tests were utilized throughout or a Friedman’s test with Dunn’s post-hoc test.

## Results

### *S. aureus* biofilm activates sustained expression of inflammatory genes in human monocytes

We followed a well-established approach (McBain 2009, Thurlow, Hanke et al. 2011, Hanke, Heim et al. 2013) to generate mature *S. aureus* biofilm in tissue culture plates, and confirmed maturation and matrix production on day 4 using staining with wheat germ agglutinin (Fig. 1A). In pilot experiments we noted that even after extensive washing of biofilm on day 4, planktonic bacteria appeared in culture supernatants after 1-3 hours, and the cultures were overgrown by *S. aureus* after 24 hours (data not shown). To avoid potential confounding of data by planktonic bacteria, we tested a strategy using different gentamycin concentrations for various lengths of time to eliminate planktonic phase *S. aureus*. We confirmed the absence of planktonic bacteria in biofilm cultures by plating supernatants on agar plates and counting CFUs and found that 50 ug/ml gentamycin for 24hrs completely suppressed planktonic *S. aureus* (Fig. 1B). In addition, 5 ug/ml gentamycin for 24 hours demonstrated very few (3 ± 1) CFUs and we used this concentration to maintain planktonic bacteria-free culture during the course of the monocyte stimulation experiments. The efficacy of suppression of bacterial outgrowth was confirmed in select experiments by plating culture supernatants on agar plates (data not shown). We added primary human monocytes with 5 ug/ml gentamycin to tissue culture plates, and harvested cells 3, 6 or 24 hrs later and measured RNA amounts using RT-qPCR (experimental design is depicted in Fig. 1C). Monocytes added to untreated tissue culture plates served as negative controls, and monocytes stimulated with the potent TLR2 agonist Pam3Cys served as positive controls. *S. aureus* biofilm strongly induced expression of prototypical inflammatory genes *IL1B*, *IL6* and TNF at the 3 hour time point where inflammatory gene induction typically peaks after stimulation of monocytes with purified TLR ligands (Ivashkiv 2011) (Fig. 1D). Biofilm induced gene expression to comparable if not greater levels than Pam3Cys (Fig. 1D and Supplementary Fig. 1). As expected, cytokine mRNA amounts returned close to baseline 24 hrs after Pam3Cys stimulation. In contrast, inflammatory cytokine gene expression remained elevated at the 24 hr time point in monocytes exposed to biofilm (Fig. 1D). These results show that *S. aureus* biofilm, in the absence of planktonic bacteria, strongly activates inflammatory genes in human monocytes in a more sustained manner than typically observed with TLR ligands.

**Figure 1.**
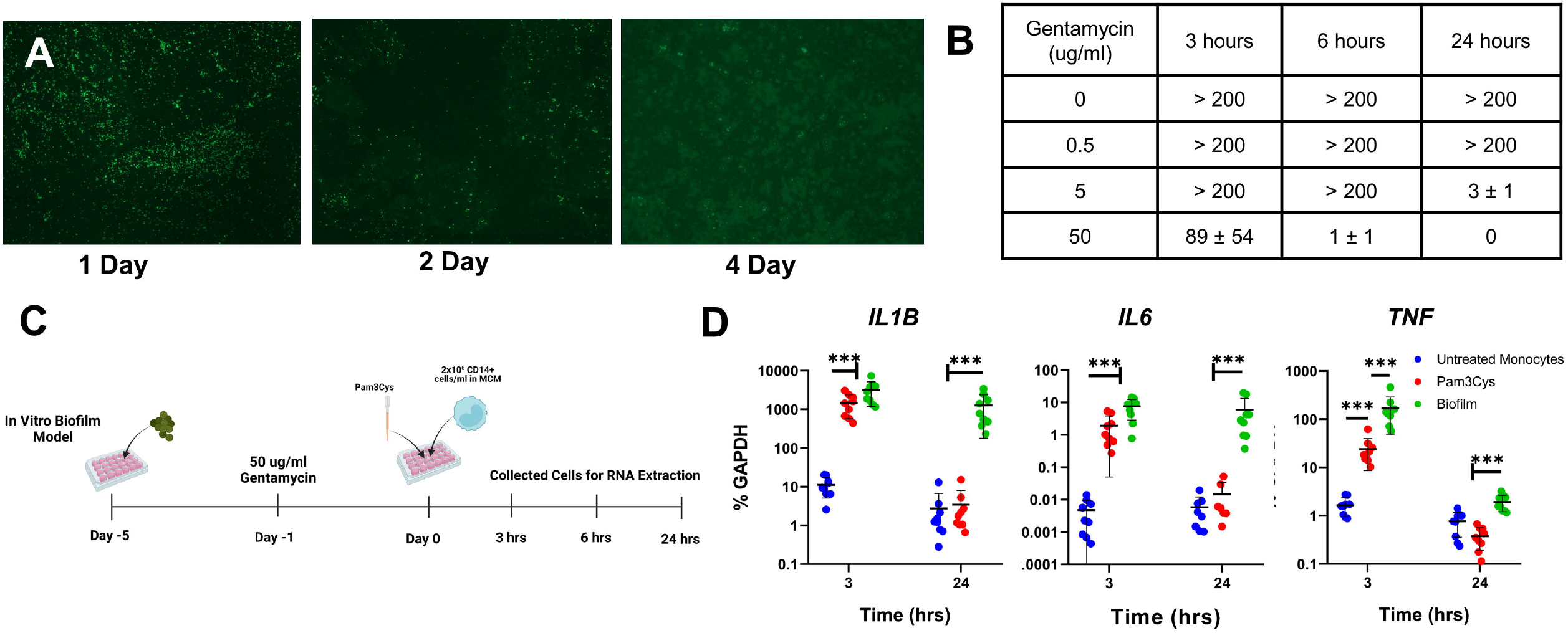
*S. aureus* biofilm induces sustained inflammatory gene expression in human monocytes. *S. aureus* biofilm was grown in tissue culture plates for 1, 2, or 4 days and stained with WGA (A). Four day mature biofilm was treated with 0, 0.5, 5, or 50 ug of Gentamycin for 3, 6 or 24hrs. Supernatant was collected and plated overnight on agar dishes and CFU were counted (n=3, B). A schematic of the biofilm culture system is presented in C. Briefly, *S. Aureus* biofilm was cultured for 4 days to mature, on the afternoon of the 4^th^ day the biofilm was treated with 50 ug/ml for 24 hrs and then purified CD14 monocytes from healthy human donors were co-cultured with the mature biofilm. *IL1B*, *IL6* and *TNF* gene expression was measured at 3 and 24 hours in 9 donors from 4 independent experiments using qPCR (D). Each dot represents one donor. Log transformed repeated measures Two-way ANOVA with Tukey’s Post-Hoc test was used, ***p<0.001.

### Direct biofilm:monocyte contact is not required for inflammatory gene induction

In the above-described experiments, monocytes could be activated by direct interaction with biofilm matrix or by soluble factors produced by biofilm-embedded bacteria. We addressed this question using a Transwell system in which biofilm is formed on the bottom of a tissue culture well, and monocytes are added to a membrane insert at the top of the well (Fig. 2A); in this system monocytes do not migrate through the membrane and do not ‘touch’ the biofilm but soluble factors in media can pass through the membrane and activate monocytes. Monocytes in the Transwell inserts were strongly activated when cocultured with biofilm, showing strong and sustained activation of *IL1B*, *IL6* and *TNF* (Fig. 2B and Supp. Fig. 2). This result suggests that *S. aureus* biofilm produces soluble factors that can activate monocytes.

**Figure 2.**
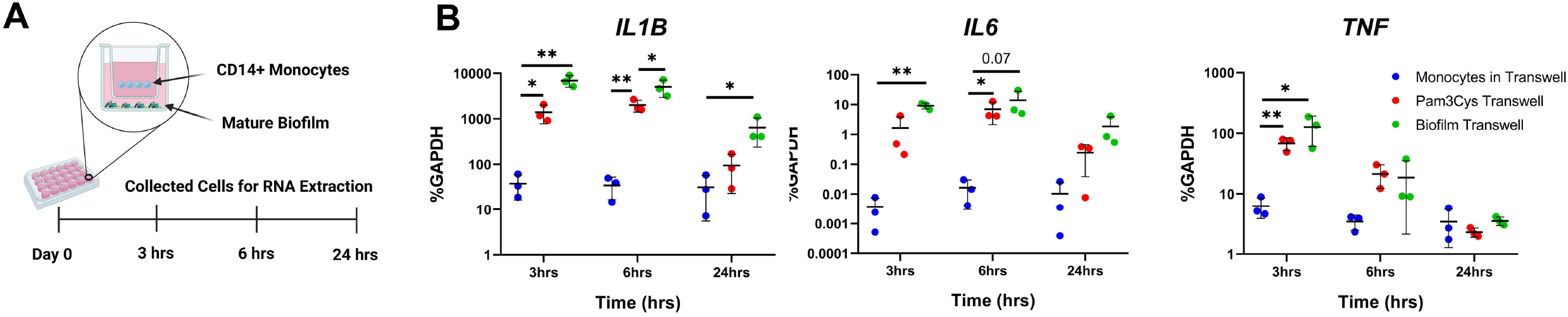
Physical contact between biofilm and monocytes is not needed to induce inflammatory cytokine genes in monocytes. *S. Aureus* biofilm was cultured for 4 days in the bottom of a Transwell plate and treated with gentamycin as previously described, then the Transwell insert was placed in the plate and CD14+ monocytes were cultured in the top of the Transwell insert (A). *IL1B*, *IL6* and *TNF* gene expression was measured at 3, 6 and 24 hours in 3 donors in 1 experiment. (B). Each dot represents one blood donor, Log transformed repeated measures Two-way ANOVA with Tukey’s Post-Hoc test, *p<0.05, **p<0.01.

### Heat-stable soluble factors produced by *S. aureus* biofilm activate monocytes

To more directly test the idea that soluble factors produced by *S. aureus* biofilm activate monocytes, we harvested supernatants generated by *S. aureus* biofilms and used them to stimulate monocytes. Biofilm supernatants effectively activated *IL1B*, *IL6* and *TNF* expression in human monocytes, strongly supporting a role for soluble factors in mediating activation (Fig. 3A and B). Bacterial biofilms secrete proteins, which are typically denatured by heat treatment, and small molecules and metabolites that are heat resistant (Yamada and Kielian 2019).

**Figure 3.**
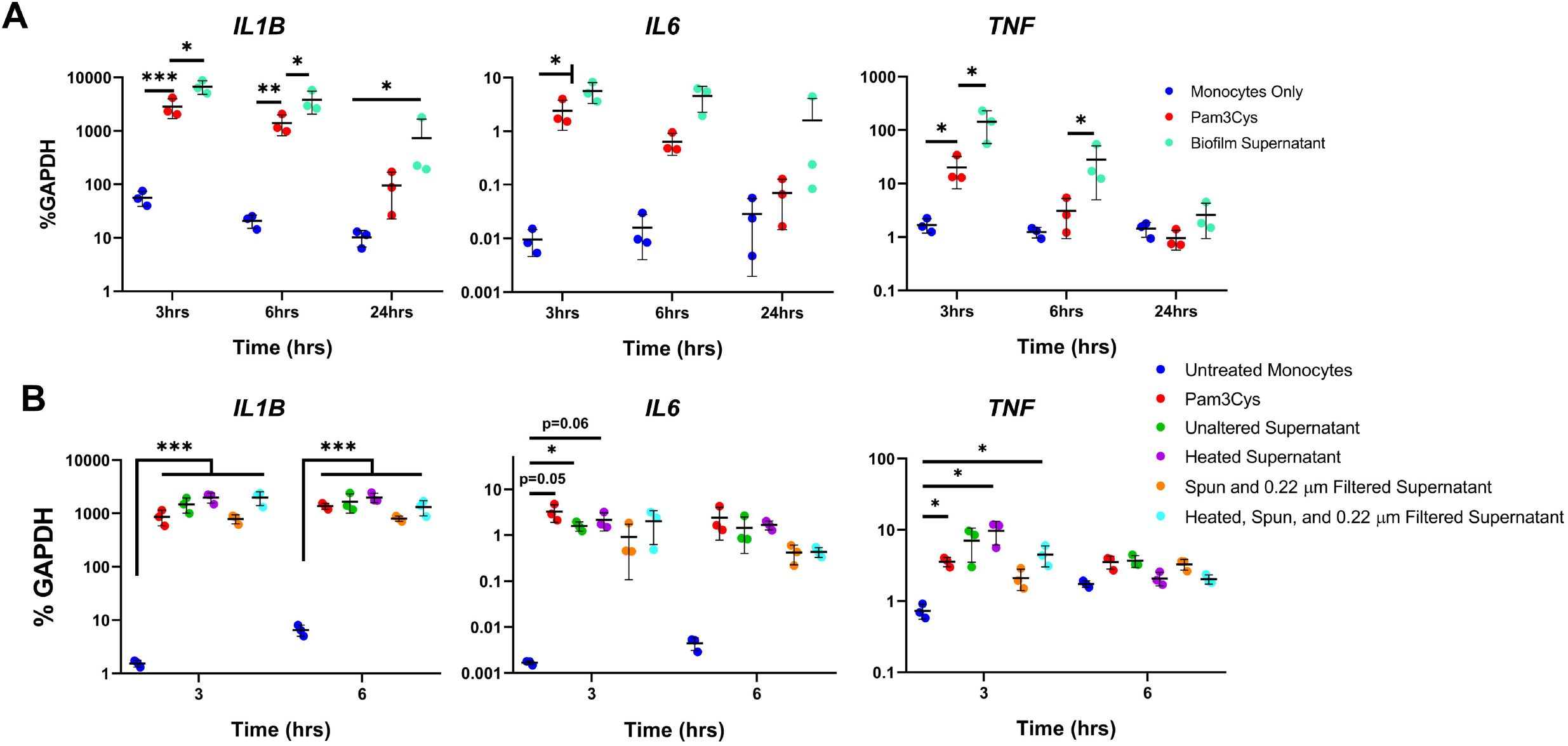
Heat-stable factors in biofilm supernatant induce inflammatory gene expression in human monocytes. *S Aureus* biofilm was cultured for 4 days in tissue culture wells and treated with Gentamycin as previously described and the supernatant was then collected. Pam3Cys or the biofilm supernatant was added to CD14+ monocytes (A). *IL1B*, *IL6* and *TNF* gene expression from was measured at 3, 6 and 24 hours in 3 donors in 1 experiment (B). Biofilm supernatant was then treated in 3 distinct ways, 1) heated to 100°C twice, 2) centrifuged at 21,000 x *g* then the supernatant was filtered at 0.22 µm, 3) the combination of both. *IL1B*, *IL6* and *TNF* gene expression was measured at 3, and 6 hours in 3 donors in 1 experiment. (C). Each dot represents one donor, Log transformed repeated measures Two-way ANOVA with Tukey’s Post-Hoc test, *p<0.05, **p<0.01.

Interestingly, biofilm supernatants preserved their activity when heated at 100°C for 30 minutes (Fig. 3B), suggesting that the bioactive factor(s) is not a protein. Supernatants also preserved their bioactivity after centrifugation and passage through a 0.22 μM filter to remove particulate debris and any residual dead bacteria (Fig. 3B). Collectively the results indicate that *S. aureus* biofilm generates heat-stable soluble factors that activate inflammatory gene expression in monocytes.

### The heat-stable activating factor(s) produced by *S. aureus* biofilm is greater than 3 kDa in size

As small molecules and metabolites are typically resistant to heat denaturation, we next used a filtration approach to test whether supernatant bioactivity was retained in a fraction containing molecules <3kDa in size. Unadulterated biofilm supernatant or supernatant that had been heated and centrifuged were passed through a 3 kDa filter and the bioactivity of the filtrate that contains <3kDa molecules was compared to the bioactivity of the original supernatants (Fig. 4A). These experiments confirmed that supernatant factor(s) that activated inflammatory gene expression were heat resistant. Strikingly, there was essentially no bioactivity in the <3 kDa fraction (Fig. 4A). The absence of viable planktonic S. aureus in these supernatants before monocyte culture (Fig. 4B) or in the culture media after 6 hrs (data not shown) was confirmed by lack of bacterial colonies after plating on agar plates. Together, the results suggest that the biofilm-derived monocyte-activating factor is heat stable and > 3 kDa in size.

**Figure 4.**
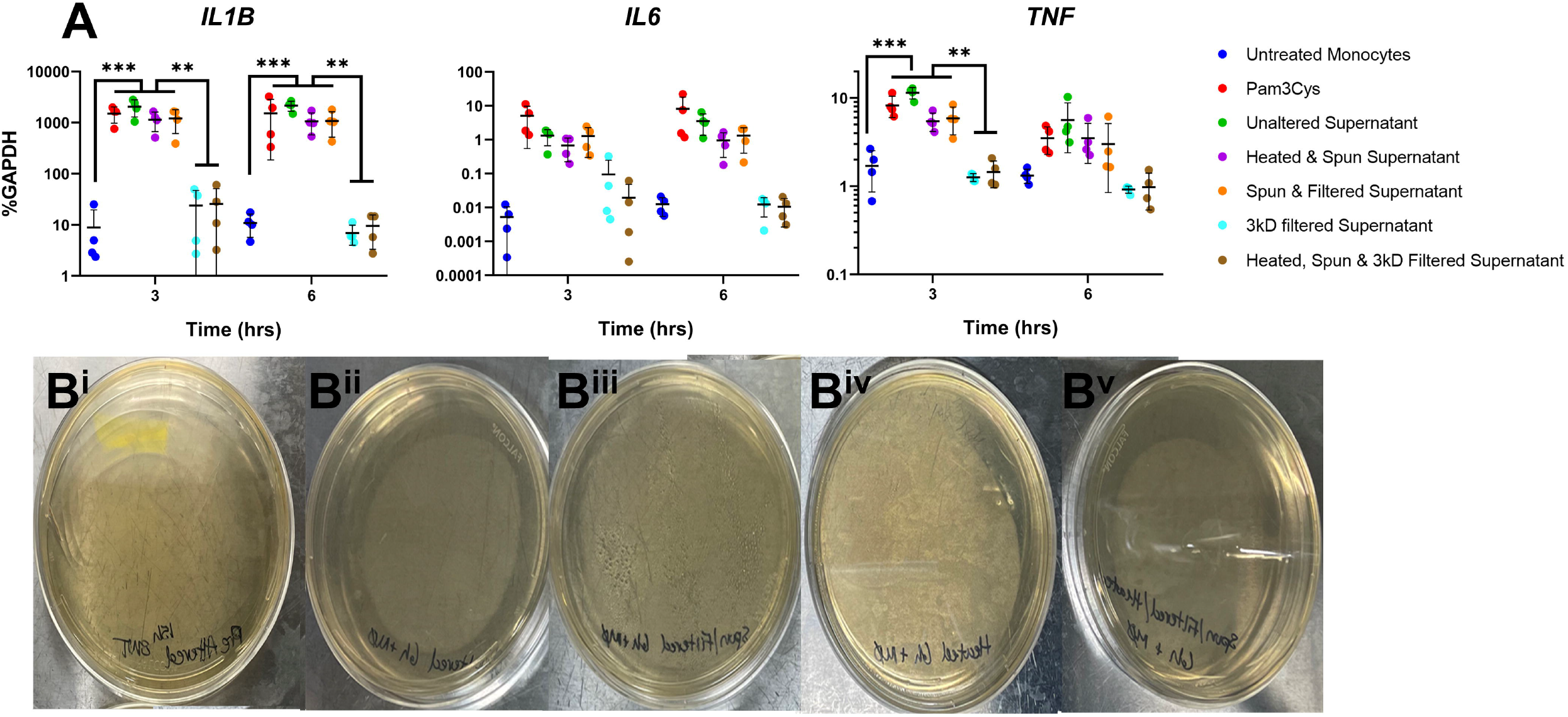
The soluble biofilm inflammatory agonist is a >3kD, non heat-denaturable molecule. Monocytes were stimulated with the various supernatant preparation for 3 and 6 hours. *IL1B*, *IL6* and *TNF* gene expression was measured from 4 donors in 1 independent experiment (A). CFU plates were generated from unaltered supernatant (B^i^); 3kD filtered (B^ii^); spun & 0.22 um filtered (B^iii^); heated (B^iv^), Spun, heated, and 3kD filtered (B^V^) before incubating with monocytes. Five CFU plates were generated for each condition and incubated at 37°C for 24 hours. Each dot represents one independent blood donor. Log transformed repeated measures Two-Way ANOVA with Tukey’s Post-Hoc test, **p<0.01, ***p<0.001.

### *S. aureus* biofilm also activates anti-inflammatory and T cell-suppressive genes in human monocytes

Activation of monocytes and macrophages by microbial products typically induces counterbalancing inflammatory and anti-inflammatory genes, to control infection while preventing over-exuberant immune responses that inflict collateral tissue damage(Ivashkiv 2011), and biofilm has been previously suggested to activate anti-inflammatory pathways and induce MDSCs (Yamada and Kielian 2019). Thus, we investigated whether *S. aureus* biofilm also activated anti-inflammatory or immune suppressive genes in human monocytes. A major negative regulator of innate immune responses (which are mediated by myeloid cells and innate lymphocytes) that has been implicated in PJI pathogenesis is the cytokine IL-10 (Fig. 5B, top panel) or supernatant (bottom panel) strongly induced expression of another suppressor of innate immunity *IL1RN*, which encodes IL-1RA, a competitive antagonist of IL-1 binding to its receptor. Suppression of T cells is a key feature of PJI (Wykes and Lewin 2018, Sokhi, Xia et al. 2022), and a major mechanism that suppresses T cells is engagement of the inhibitory PD-1 receptor by PD-1 ligands that can be expressed on myeloid cells (McLane, Abdel-Hakeem et al. 2019, Chamoto, Yaguchi et al. 2023, Sharma, Goswami et al. 2023). Strikingly, biofilm and its supernatant strongly induced expression of both PD-1 ligands PD-1L1 (CD274) and PD-1L2 (CD273) (Figure 5C). Biofilm induced these negative regulators more strongly than Pam3Cys at most time points (Fig. 5A – 5C). Biofilm also induced expression of MDSC marker genes (Veglia, Sanseviero et al. 2021) such as *IDO1*, *CEBPB*, *PTGS2*, *S110A8* and *S100A9*, which may contribute to the T cell suppression observed in PJI (Fig. 5D). These results show that *S. aureus* biofilm strongly induces anti-inflammatory and T cell-suppressive genes, and suggest that biofilm can activate these suppressive genes more strongly than a TLR ligand like Pam3Cys.

**Figure 5.**
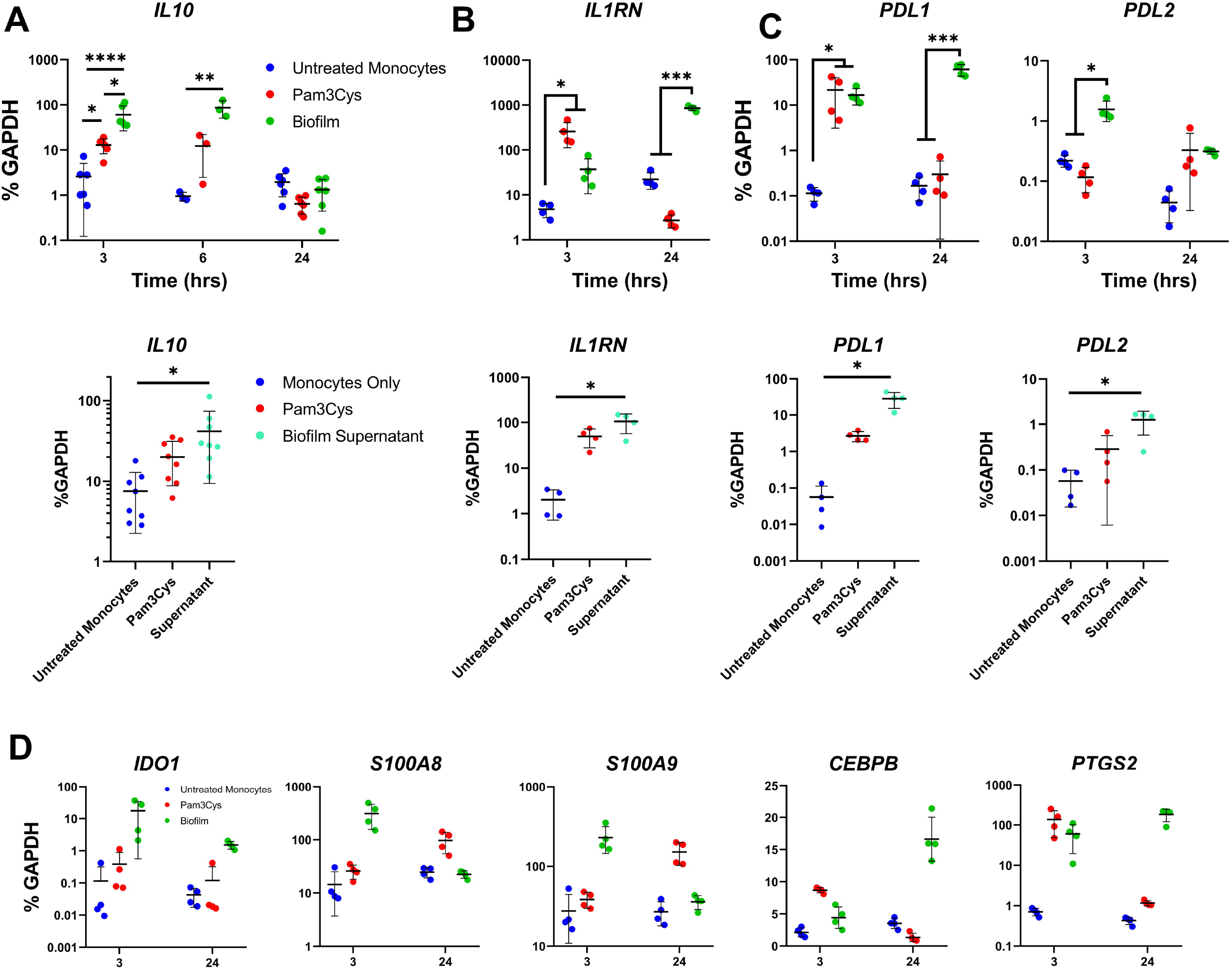
*S. aureus* biofilm and supernatant activate anti-inflammatory and immune-suppressive genes. CD14+ monocytes were stimulated with Pam3Cys, biofilm supernatant or day 4 matured *S. Aureus* biofilm for 3, 6 or 24 hrs. Gene expression of *IL10* (A), *IL1RN* (B), *PDL1* or *PDL2* (C), *IDO1*, *S100A8*, *S100A9*, *CEBPB* and *PTGS2* (D). Biofilm experiments were performed in 3-6 donors in 1-2 independent experiments (same donors as Fig 1). Supernatant experiments were performed in 4-8 donors in 1-2 independent experiments. Each dot represents one independent blood donor, log transformed repeated measures Two-way ANOVA with Tukey’s Post-Hoc test (A-C, top; D), Friedman’s Test with Dunn’s Post-Hoc (A-C, bottom), *p<0.05, **p<0.01, ***p<0.001.

### Jak signaling regulates biofilm-mediated gene induction

Signaling pathways that drive expression of the above-described inhibitory genes are not well characterized. Stimulation of monocytes by bacterial products typically activates Jak-STAT signaling by autocrine-produced cytokines. Baricitinib, an inhibitor of Jak1 and Jak2 kinases, significantly suppressed biofilm-mediated induction of *IL1RN* and *PD1L1* at 3 hours (Fig 6 and Supplemental Fig. 4). In contrast, Jak inhibition had minimal effects on TNF expression. These results suggest that Jak inhibition may be an effective therapeutic strategy to attenuate biofilm-induced inhibitory gene expression.

**Figure 6.**
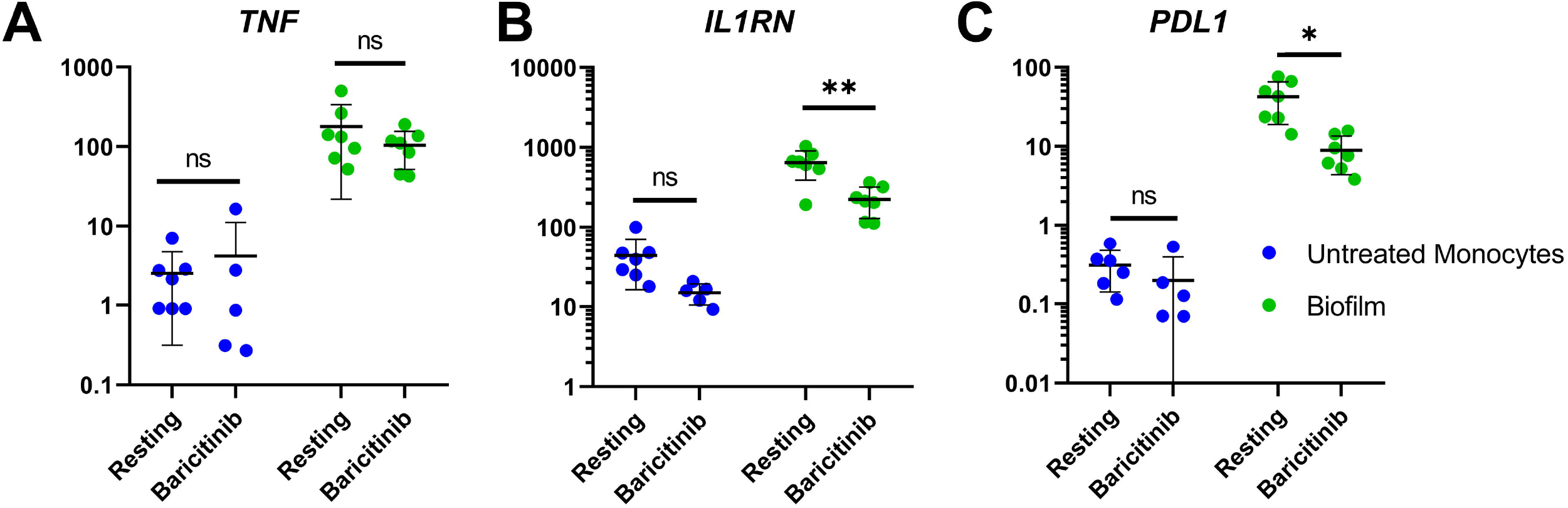
Jak inhibition reduces induction of T cell-suppressive genes by biofilm. Baricitinib was incubated with monocytes plus biofilm for 3 hours in 5-7 donors over 3 independent experiments. Gene expression was measured for *TNF* (A), *IL1RN* (B) and *PDL1* (B). Each dot represents one independent blood donor, log transformed repeated measures Two-Way ANOVA with Tukey’s Post-Hoc test, *p<0.05.

## Discussion

A paradigm in the orthopedic PJI field is that biofilm acts predominantly as a shield to protect and sequester bacteria from immune cells, thereby providing a reservoir of bacteria that enters the planktonic phase and fuels persistent infection (Scherr, Heim et al. 2014, Masters, Ricciardi et al. 2022). The data presented in this manuscript supports the emerging evolution of this paradigm to include a more active role for biofilm in modulating immune responses, and show that this modulation is mediated by soluble factors secreted by *S. aureus* biofilm. To our knowledge, these studies represent the first attempt to rigorously exclude planktonic bacteria from the analysis of biofilm-immune cell interactions. *S. aureus* biofilm induced expression of both inflammatory and immune suppressive genes in human monocytes in vitro, which is in line with expression of these genes at the biofilm interface in a mouse PJI model(Farnsworth, Schott et al. 2018, Tomizawa, Ishikawa et al. 2020, Sokhi, Xia et al. 2022) and with inflammatory infiltrates in human PJI (Masters, Bhagwate et al. 2022). Induction of suppressive genes by biofilm was sensitive to suppression by Jak inhibition. These results reveal a dichotomous activation of human innate immune cells by soluble biofilm products and provide insight into regulation of suppressive monocyte genes via Jak-STAT signaling.

An emerging idea is that biofilm is not solely a physical barrier but a living tissue that can affect immune cells via production of small molecule metabolites or proteins toxins (Scherr, Heim et al. 2014, Scherr, Hanke et al. 2015, Yamada and Kielian 2019). Our work supports this idea, but surprisingly the major monocyte-activating bioactivity in biofilm supernatants was not a small molecule and very unlikely a protein, as it was resistant to heat denaturation. Bacterial components that activate innate immune cells are termed pathogen-associated molecular patterns (PAMPs) (Pidwill, Gibson et al. 2020) and heat-resistant PAMPs include cell wall components including glycans, and nucleic acids. Another interesting possibility is that biofilm-enmeshed *S. aureus* actively secrete DNA-containing microvessicles that can activate monocytes, as has been described for microbiota (Erttmann, Swacha et al. 2022), and would not have been removed during processing of the supernatants. However, nucleic acids would be expected to induce a strong IFN response in monocytes, which we did not observe, and pilot experiments using RNAse and DNase showed no effect. Thus, we favor that biofilm bacteria are producing either large glycan/polysaccharide complexes or shedding cell wall fragments. Interestingly, production of such activating factors by living bacteria would classify them as a ‘vita-PAMP’ that signals microbial viability to alert the immune system (Blander and Barbet 2018). Identification of these immune-activating factors will require extensive biochemical fractionation and purification in future work.

An important question relates to the balance between induction of pro-inflammatory versus suppressive genes in monocytes by *S. aureus* biofilm-derived factors. Our in vitro system showing strong activation of canonical inflammatory genes such as *TNF*, *IL6* and *IL1B* is in accord with previous work in vitro and in vivo in human PJI and mouse models (Krishna and Miller 2012, Menousek, Horn et al. 2022, Sokhi, Xia et al. 2022). Although the innate immune cytokines TNF and IL-1β have been implicated in control of *S. aureus* infections, strong activation of inflammatory cytokines and innate immune cells seems insufficient to clear *S. aureus* biofilm-associated infections(Miller and Cho 2011, Krishna and Miller 2012, Alphonse, Rubens et al. 2021, Sokhi, Xia et al. 2022). One possible explanation for such lack of clearance is attenuation of cytokine production by IL-10, which was induced in our system and has been previously suggested to promote PJI based on in vitro and in vivo data (Heim, Vidlak et al. 2014, Heim, Vidlak et al. 2015, Heim, Vidlak et al. 2015, Tebartz, Horst et al. 2015, Heim, Vidlak et al. 2018, Heim, Bosch et al. 2020, Sokhi, Xia et al. 2022). We additionally describe strong activation of *IL1RN*, which would contribute to dampening innate immune responses by suppressing IL-1 signaling. Another plausible explanation for lack of clearance of *S. aureus* infections is suppression of adaptive immunity and T cell responses. Although suppression of T cells in orthopedic PJI has been described and attributed to MDSCs (Tebartz, Horst et al. 2015, Heim, Vidlak et al. 2018, Sokhi, Xia et al. 2022), mechanisms of T cell suppression are not known. By blocking IL-1β interaction with its receptor, IL-1RA would also suppress the differentiation of Th17 T cells important for anti-Staph immunity. Our study also shows that *S. aureus* biofilm strongly induces expression of PD-1 ligands, which are potent suppressors of T cells and implicated in immunoparalysis that occurs in tumors. Thus our work supports a model whereby dichotomous induction of inflammatory and suppressive factors by biofilm holds inflammation in check to limit tissue damage, while suppressing T cell immunity to prevent clearance of infection. This model suggests that therapeutic targeting of PD-1L – PD-1 interactions may increase the effectiveness of anti-Staph immunity, similarly to what has been observed by induction of anti-tumor immunity by immune checkpoint blockade that targets this pathway.

An alternative and complementary therapeutic strategy would be to target inhibitory cytokines and signaling pathways activated by biofilm. IL-10 is well known to inhibit production of inflammatory cytokines, but its role in expression of anti-inflammatory cytokines (like IL-1RA) and T cell-suppressive molecules (like PD-1 ligands) is not well characterized. Our attempts to block autocrine IL-10 function in our system using IL-10 neutralizing and IL-10 receptor blocking antibodies have yielded inconsistent results to date, possible secondary to inefficiency of IL-10 blockade. We then took the alternative strategy of inhibiting Jaks, which clearly implicated Jak signaling in the induction of IL1RA and PD-1 ligands. Although myeloid cell receptors that are activated by *S. aureus* PAMPs such as TLR2 and Dectin-1 that do not typically activate a strong IFN response, which is in accord with our data, interpretation of the Jak inhibitor data is subject to the caveat that Jak inhibition will also block autocrine signaling by cytokines other than IL-10 in our system.

The limitations of our study are mainly related to using an in vitro system to model *S. aureus* biofilm effects on human monocytes. Specifically, we utilized a static flow system with culture plastic to grow our biofilm which may not fully re-capitulate the in vivo, metal or the bone growth environment (McBain 2009). Although our results recapitulate previous in vivo findings in human PJI and mouse models (Carli, Ross et al. 2016, Carli, Bhimani et al. 2017, Sokhi, Xia et al. 2022) it will be important to more thoroughly investigate in vivo our novel results on induction of inhibitory molecules and pathways. We also only partially characterized the biofilm-produced factor(s) that activates monocytes and future biochemical experiments will be required to identify the vita-PAMP that is produced by *S. aureus* biofilm to activate human monocytes.

## Figure Legends

**Supplemental Figure 1. An additional set of 3 donors to corroborate Figure 1.** *IL1B*, *IL6* and *TNF* gene expression was measured at 3, 6 and 24 hours in 3 additional donors in 1 experiment. Each dot represents one donor. Log transformed repeated measures Two-way ANOVA with Tukey’s Post-Hoc test was used, ***p<0.001.

**Supplemental Figure 2. An additional set of 3 donors to corroborate Figure 2.** *S Aureus* biofilm was cultured for 4 days in the bottom of a Transwell plate and treated with gentamycin as previously described then Transwell inserts were placed in the wells and CD14+ monocytes were cultured in the top of the Transwell. *IL1B*, *IL6* and *TNF* gene expression was measured at 3, 6 and 24 hours in 3 donors in 1 experiment. (B). Each dot represents one donor, Log transformed repeated measures Two-way ANOVA with Tukey’s Post-Hoc test, *p<0.05, **p<0.01.

**Supplemental Figure 3. An additional set of 5 donors to corroborate Fig. 3. Biofilm supernatant induces a sustained IL-1β response in human monocytes.** *S Aureus* biofilm was cultured for 4 days tissue plates and treated with Gentamycin as previously described and the supernatant was then collected. Pam3Cys or the biofilm supernatant was cultured with CD14+ monocytes. *IL1B*, *IL6* and *TNF* gene expression from was measured at 3 and 6 hr in 5 donors (except for heated, spun and filtered condition where n=2, this group was omitted from statistical analysis) from 2 independent experiments (A). Each dot represents one donor, Log transformed repeated measures Two-way ANOVA with Tukey’s Post-Hoc test, *p<0.05, **p<0.01.

**Supplemental Figure 4. An additional set of 3 donors and Pam3Cys control to corroborate Fig. 6. Jak inhibition reduces induction of T cell-suppressive genes by biofilm.** Baricitinib was incubated with monocytes for 1hr before stimulating with either Pam3Cys or biofilm for 3 hours. Gene expression was measured for *TNF* (A), *IL1RN* (B) and *PDL1* (C). Log transformed repeated measures Two-Way ANOVA with Tukey’s Post-Hoc test, *p<0.05. n=3 per group.

**Supplemental Table 1.**
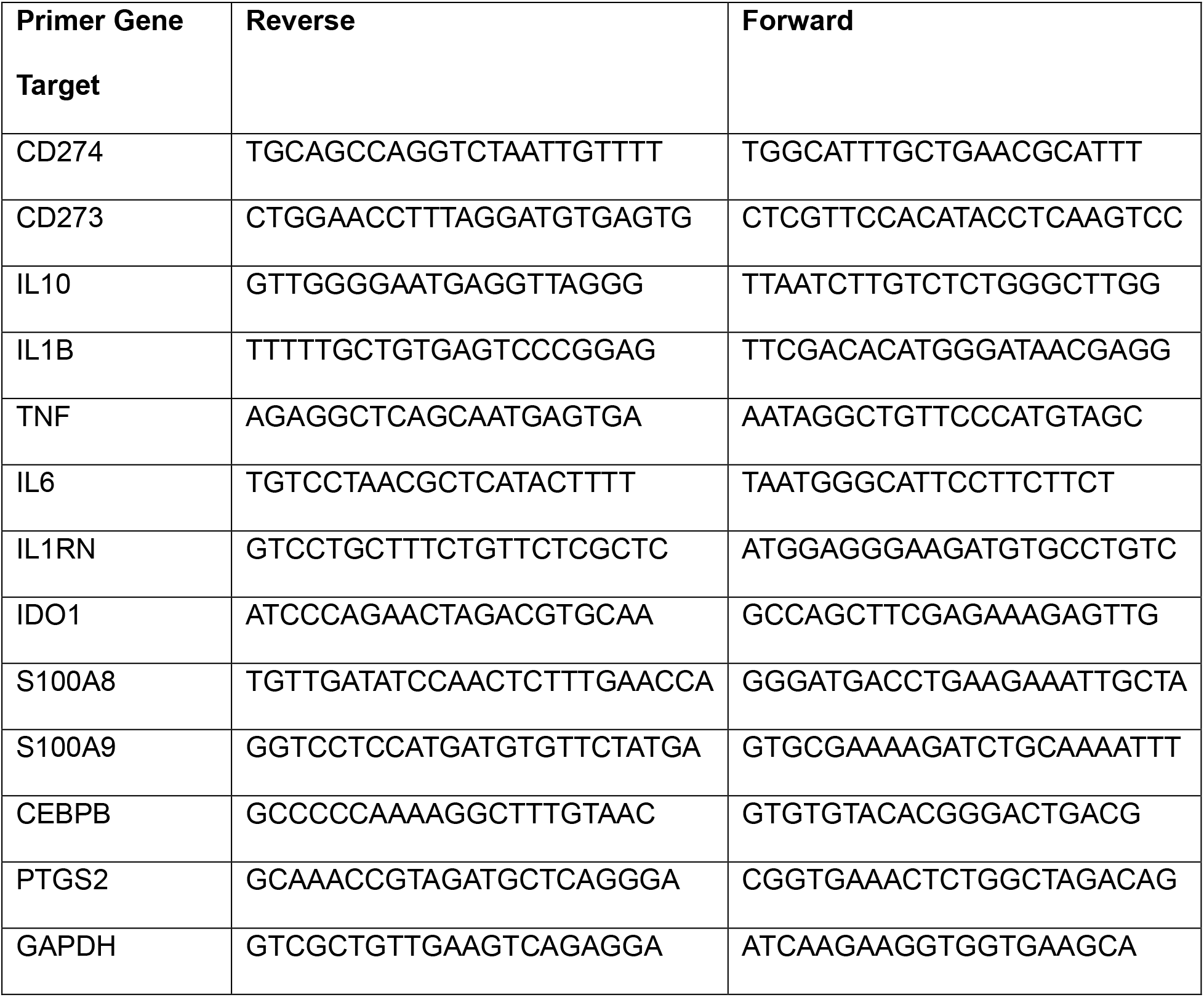
Primers used for qPCR.

