## Supplementary figures and images for "*Staphyloccocus aureus* biofilm, in the absence of planktonic bacteria, produces factors that activate counterbalancing inflammatory and immune-suppressive genes in human monocytes"

### Supplemental Figure 1

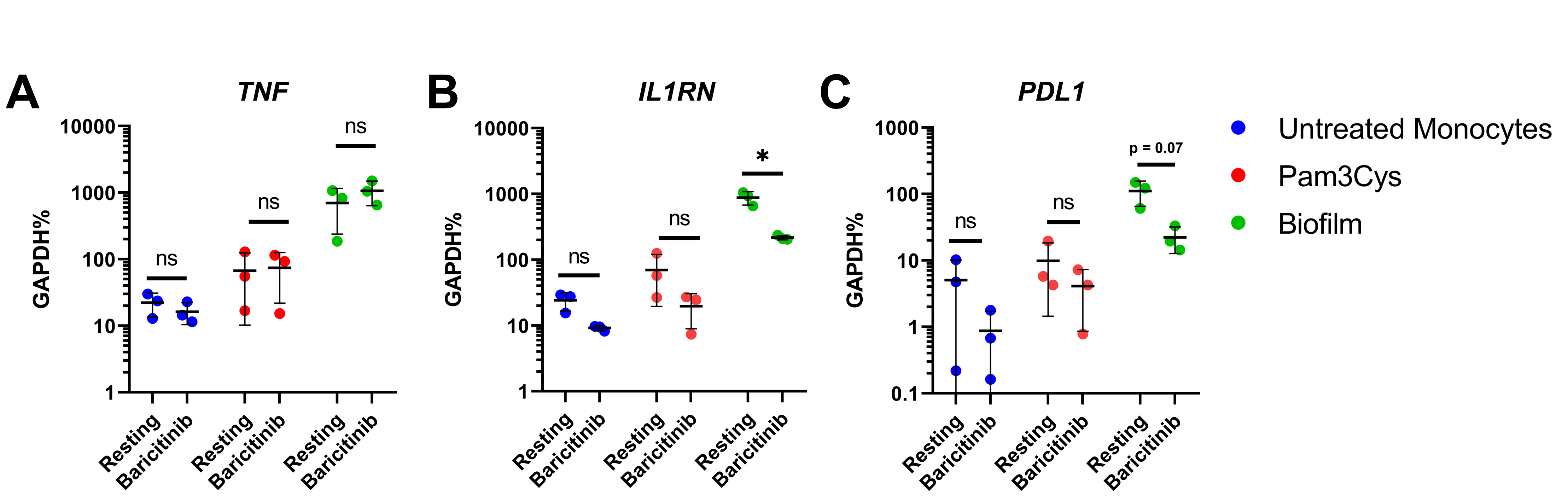

### Supplemental Figure 2

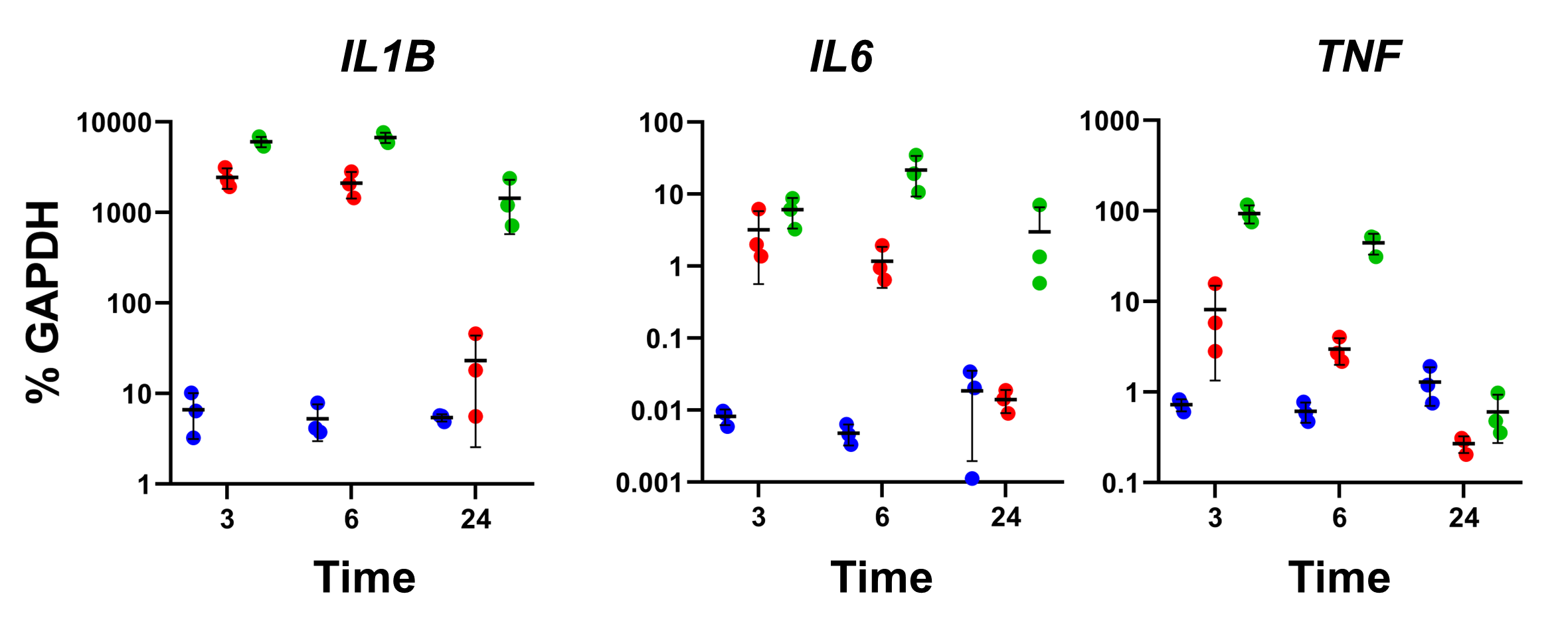

### Supplemental Figure 3

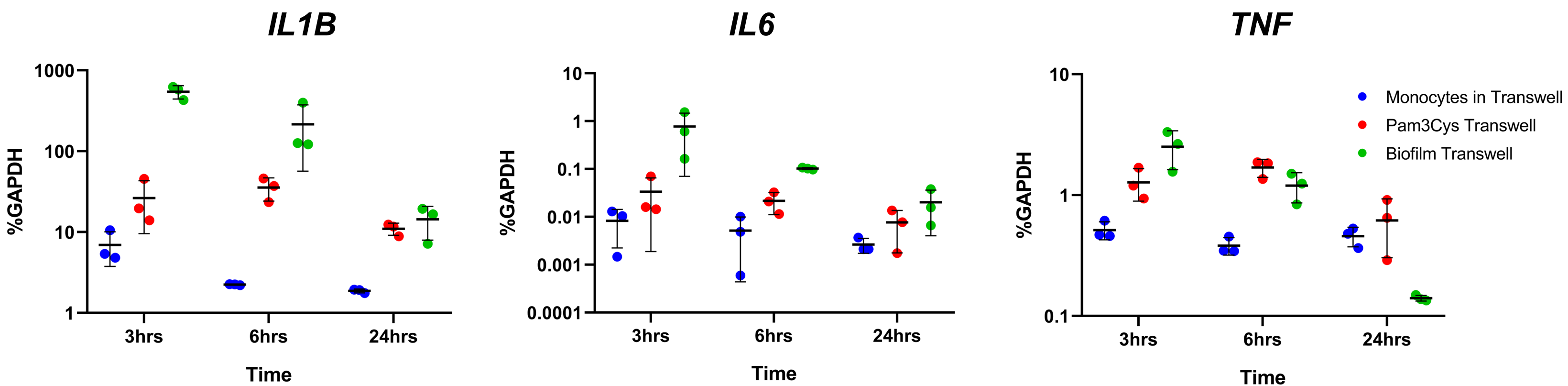

### Supplemental Figure 4

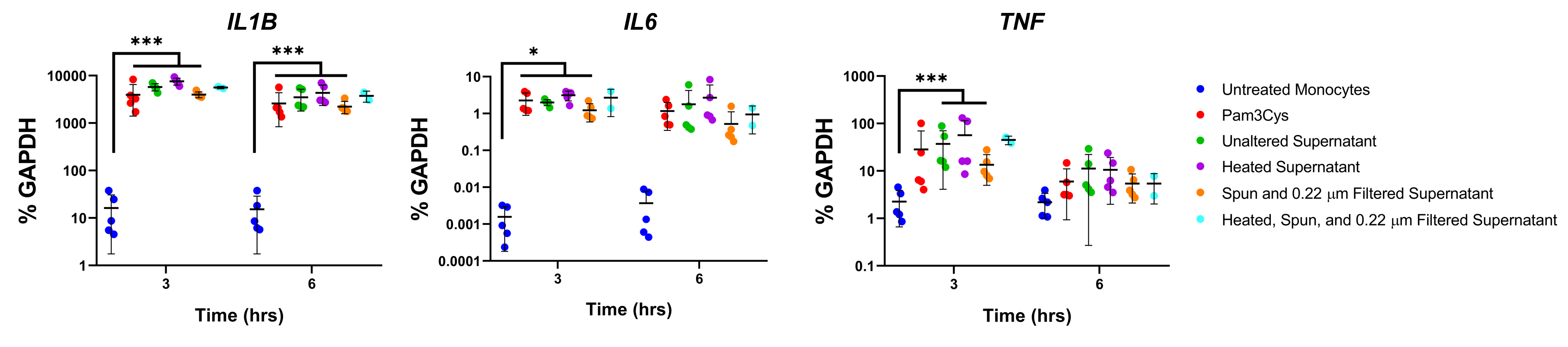
